# The challenge of heterogeneous evidence in conservation

**DOI:** 10.1101/797639

**Authors:** Alec P. Christie, Tatsuya Amano, Philip A. Martin, Silviu O. Petrovan, Gorm E. Shackelford, Benno I. Simmons, Rebecca K. Smith, David R. Williams, Claire F. R. Wordley, William J. Sutherland

## Abstract

Conservation efforts to tackle the current biodiversity crisis need to be as efficient and effective as possible. To inform decision-makers of the most effective conservation actions, it is important to identify biases and gaps in the conservation literature to prioritize future evidence generation. We assessed the state of this global literature base using the Conservation Evidence database, a comprehensive collection of quantitative tests of conservation actions (interventions) from the published literature. For amphibians and birds, we investigated the nature of Conservation Evidence spatially and taxonomically, as well as by biome, effectiveness metrics, and study design. Studies were heavily concentrated in Western Europe and North America for birds and particularly amphibians. Studies that used the most robust study designs - Before-After Control-Impact and Randomized Controlled Trials - were also the most geographically restricted. Furthermore, there was no relationship between the number of studies in each 1×1 degree grid cell and the number of species, threatened species or data-deficient species. Taxonomic biases and gaps were apparent for amphibians and birds - some orders were absent from the evidence base and others were poorly represented relative to the proportion of threatened species they contained. Temperate forest and grassland biomes were highly represented, which reinforced observed geographic biases. Various metrics were used to evaluate the effectiveness of a given conservation action, potentially making studies less directly comparable and evidence synthesis more difficult. We also found that the least robust study designs were the most commonly used; studies using robust designs were scarce. Future research should prioritize testing conservation actions on threatened species outside of Western Europe and North America. Standardizing metrics and improving the robustness of study designs used to test conservation actions would also improve the quality of the evidence base for synthesis and decision-making.

## Introduction

Biodiversity conservation does not receive sufficient funding to effectively combat the biodiversity crisis (Dirzo et al. 2014). This means that conservation researchers must prioritize research effort to maximize its potential to inform conservation efforts. Knowing the current state of the evidence base for conservation is crucial to prioritizing future research efforts (Aranda et al. 2011). Whilst evidence-based conservation is ultimately likely to lead to more efficient conservation efforts, this approach requires a stable and reliable evidence base (Koricheva & Kulinskaya, 2019). Efforts to summarize the evidence in conservation relating to the effectiveness of different conservation actions (‘interventions’; Sutherland et al. 2004) have produced a substantial evidence base (Sutherland et al. 2019), yet little is known about the biases, gaps and clusters of this evidence. In this paper, we focus on studies that test conservation interventions, such as restoring or creating grasslands for birds or creating ponds for amphibians.

The lack of resources in conservation are likely to lead to several forms of heterogeneity in the evidence base for conservation. These forms of heterogeneity may determine our ability to provide relevant evidence-based recommendations to decision-makers, or make the process of evidence synthesis more challenging. For example, geographical and taxonomic biases towards certain regions or groups (e.g. North America and Europe or charismatic species) may lead to little evidence being available for certain local contexts. Alternatively, heterogeneity could be useful if research effort is prioritized to where it is needed most in conservation - for example, more studies on threatened species than non-threatened species. Patterns in research effort are also influenced by the physical accessibility of locations to researchers from the Global North (Reddy and Dávalos 2003) and multiple socio-economic variables (e.g. GDP per capita, affluence, language, security, conflict and infrastructure; Fisher et al. 2011, Martin, Blossey, and Ellis 2012; Amano and Sutherland 2013; Meyer et al. 2016; Hickisch et al. 2019). These factors are likely to affect the representation of different biomes or habitats in the evidence base (Fazey et al. 2005). Research effort is also known to vary with taxonomic group (Clark & May 2002; Murray et al. 2015; Donaldson et al. 2016), and to depend on the range size, diet and body size of species (Brooke et al. 2014), contributing to biases towards larger, more detectable species (Brodie 2009; Cardoso et al. 2011). These forms of heterogeneity affect the external validity of studies in the evidence base and are therefore important to help us understand how much evidence is available to inform conservation in different contexts.

Other forms of heterogeneity may instead complicate the synthesis of evidence. Heterogeneity in the usage of different types of metrics to assess the effectiveness of the same conservation intervention may make approaches, such as meta-analyses, difficult to use. This is because studies are less directly comparable if they use different metrics to assess effectiveness, thus reducing the number of studies that can be combined in a given meta-analysis. Clearly, different metrics may be useful to assess different aspects of an intervention’s effectiveness. However, wide variation in metrics used to test the same intervention could cause confusion for decision-makers, especially if studies using different metrics yield different results (Mace & Baillie 2007; Capmourteres & Anand 2016).

Differences in study quality, for example due to the usage of different study designs, may also make it more difficult to decide which studies to trust over others, particularly if they give conflicting results. Several different study designs are used to assess impacts of threats and interventions in ecology (De Palma et al. 2018; Christie et al. 2019), all of which are affected by different sources and levels of bias and noise. These range from relatively robust designs such as experimental Randomized Controlled Trials (RCTs) and quasi-experimental Before-After Control-Impact designs (BACI), to less robust designs such as Control-Impact (CI), Before-After (BA) and After (also called time series). Evidence may also come in the form of systematic reviews and meta-analyses, generally considered robust depending on their methodology and the robustness of the studies they include. Typically, the evidence base for conservation is considered to have relatively few studies with robust study designs, due to logistical, funding, and time-based constraints (De Palma et al. 2018; Christie et al. 2019). However, we do not know how this broad pattern varies geographically, or how this translates into the prevalence of different designs at an intervention scale. Insufficient reliable evidence would mandate greater efforts to improve the type of study designs we use.

The aim of this study is to improve our empirical and quantitative understanding of the heterogeneity, biases and knowledge gaps in the evidence base for conservation. To do this, we present a series of analyses of the Conservation Evidence database (Sutherland et al. 2019), a comprehensive collection of 5494 publications (as of September 2019) that have quantitatively tested the effectiveness of conservation interventions. To quantify heterogeneity in this evidence base we set out to answer several research questions: 1) what is the geographic distribution of studies?; 2) how does this distribution vary for studies with different designs?; 3) what is the taxonomic distribution of studies?; and for studies on a given conservation intervention, how much variation is there in the use of 4) different study designs, and 5) different metrics? Identifying patterns, biases, and knowledge gaps in the evidence base can help in setting priorities for future research. With a more robust and more complete evidence base, research can better support evidence-based decision making in conservation and ultimately more effective biodiversity conservation.

## Methods

### Conservation Evidence database

The Conservation Evidence project summarizes studies that have quantitatively tested the effect of a conservation intervention - defined as any management practice or action that a practitioner may undertake to benefit biodiversity (Sutherland et al. 2019). These studies are found using systematic manual searches of the conservation literature, including over 290 English and 150 non-English language journals (Sutherland et al. 2019). The Conservation Evidence website is structured into over 1,870 different interventions (e.g. control invasive mammals on islands) contained within 15 synopses (e.g. Bird Conservation) and displays a summary of each study included, or multiple summaries if a study’s results apply to several interventions (e.g. both pond creation and translocation of amphibians). A list of interventions is created for each synopsis through consulting initial literature scans and an advisory board (a mix of academics, practitioners and policymakers with subject-specific expertise; Sutherland et al. 2019). Interventions are usually described at a fine scale (for example, “Set longlines at the side of the boat to reduce seabird bycatch” is a separate intervention from “Set lines underwater to reduce seabird bycatch”).

As we wanted to assess the number of studies per intervention for different subsets of studies (e.g. taxa or biome), we grouped some interventions that were essentially the same type of intervention but focused on single taxa or habitats (e.g. “create ponds for frogs” and “create ponds for toads” would be grouped into “create ponds”; see Appendix S1). This ensured that the scope of interventions was appropriate for our analysis and that we did not artificially reduce the mean number of studies per intervention.

We extracted metadata from the database of the website for every study within the amphibian (n=410; Smith & Sutherland 2014) and bird synopses (n=1,239; Williams et al. 2012), including the latitude and longitude coordinates (mean coordinates where a study used multiple sites). We only considered studies for amphibians and birds as these taxa had the most complete and comprehensive metadata in the database. The searches that retrieved these studies from the literature (see Sutherland et al. 2019) were last conducted in 2012 for amphibians and 2011 for birds. Whilst these searches are not as recent as we might wish, these data provide the only way to reasonably assess biases in a large number of studies that have tested the effectiveness of conservation interventions. For all analyses that quantify the mean number of studies per intervention we excluded interventions that originally did not contain any studies (i.e. no studies present regardless of biome, metric or design types; 31 interventions for Amphibians and 56 for birds as of September 2019).

### Patterns in evidence for different metrics and designs

A standardized set of keywords are used to describe study design in the Conservation Evidence database, which we used to classify designs (Table 1). To determine the accuracy of reported study designs, we manually checked the original papers of a random 5% of studies in the database (n=21 for Amphibians; n=62 for Birds). The correct design was assigned to 95% of amphibian studies and 94% of bird studies (see Appendix S2). A single study in the Conservation Evidence database can have multiple designs, if there are multiple parts to a study published in the same report or paper; when a single study has several different designs, we counted each separately. An individual study can also be assigned to multiple interventions and multiple synopses if it contains relevant information. We used the number of studies with each design per intervention as the major variable of interest.

**Table 1.** Definitions for each study design based on the criteria used to define them, and the keywords used, in the Conservation Evidence database (Sutherland et al. 2019).

| Design | Controlled? | Sampling before intervention? | Randomized allocation of intervention and control? |
| --- | --- | --- | --- |
| Keywords used | “Controlled” or “Site comparison”? | “Before-and-After” OR “Before-After”? | “Randomized”? |
| After | No | No | No |
| BA | No | Yes | No |
| CI | Yes | No | No |
| RCT | Yes | No | Yes |
| BACI | Yes | Yes | No |

For metrics measuring effectiveness of interventions, we put together and tested a set of regular expression rules (e.g. matching keywords and patterns) to detect the following from text extracted via web scraping: abundance, density, cover, diversity/species richness, reproductive success, mortality and survival (Appendix S3). We aggregated abundance, density and cover into one group, and mortality and survival into another, creating four metric types (abundance/density/cover, diversity/species richness, mortality/survival and reproductive success). This allowed us to investigate the number of studies per intervention for each metric type.

For web scraping we used the XML package (Lang and CRAN team 2018a) and RCurl package (Lang and CRAN team 2018b) in R statistical software version 3.5.1 (R Core Team 2018). We also used the doParallel package (Microsoft Corporation & Weston 2017) to increase computational performance. For a random 5% of studies (n=21 amphibians, n=62 birds) we found that the metrics extracted by web scraping were correct for 81% of amphibian studies and 82% of bird studies (Appendix S4). While automating the extraction of study design and effectiveness metrics means a small proportion of studies were misclassified, our method offers the most feasible and reproducible methodology to analyze the entire evidence base and controls for some potential biases that would affect manual classification (Stouffer et al. 2014).

### Patterns in evidence spatially and taxonomically

We mapped the spatial distribution of studies in the database by creating a raster layer with the raster package (Hijmans 2019), where the number of studies was summed for each 4×4 degree cell using longitude and latitude coordinates – we used a 4×4 degree resolution to aid data visualization for the maps we produced. We excluded reviews from our analyses as they were often global or regional in scale. To estimate the geographical coverage of studies we counted the number of countries and continents they were present in. We also compared the number of studies in each 1×1 degree cell with the number of species, threatened species and data-deficient species for extant amphibian and bird species using data downloaded from the International Union for Conservation of Nature (IUCN) Red List (IUCN 2019). We attempted to quantify the relationship between these variables by using generalized linear models with a quasi-Poisson error distribution, as well as zero-inflated Poisson and negative binomial models, but model assumptions (in terms of overdispersion, normally distributed residuals and patterns between residuals and fitted values) were still violated. Therefore, we decided to use the infotheo package (Meyer 2014) to assess the mutual information value between the number of studies and the number of species, threatened species and data-deficient species (higher values indicate more similar distributions, a value of zero indicates the distributions are independent of each other). This approach gave us an indication of the amount of information obtained about the number of studies from observing the number of species, threatened species and data-deficient species (i.e. how similar were the geographical distributions of studies and species?) without violating assumptions of generalized linear models.

We also used the coordinates for each study and a shapefile of terrestrial biomes (Dinerstein et al. 2017) to assign studies to a biome (sp package in R; Pebesma & Bivand 2005; Bivand et al. 2013). We assigned studies that did not fall within terrestrial biomes to a biome termed “Non mangrove-marine” (e.g. studies considering seabirds over ocean or on cliffs of remote islands). Terrestrial biome polygons were used for the following biome types: boreal forests/taiga; deserts & xeric shrublands; flooded grasslands & savannas; Mediterranean forests, woodlands & scrub; montane grasslands & shrublands; temperate broadleaf & mixed forests; temperate conifer forests; temperate grasslands, savannas & shrublands; tropical & subtropical coniferous forests; tropical & subtropical dry broadleaf forests; tropical & subtropical grasslands, savannas & shrublands; tropical & subtropical moist broadleaf forests; tundra and mangroves. We then calculated the mean number of studies per intervention for each biome. We also calculated the mean number of studies per intervention for studies from any forest biome and any grassland or shrubland biome. Similarly, we also calculated the mean number of studies with each type of study design and metric per intervention.

For investigating the distribution of evidence taxonomically, we considered the major bird orders according to the IUCN (2019) Red List and a cladogram based on Prum et al. (2015). For amphibians, we considered the three major orders based on the IUCN Red List and a trimmed cladogram from Pyron & Wiens (2011). We obtained data on the number of species and threatened species (with vulnerable, endangered or critically endangered status) from the IUCN Red List website (https://www.iucnredlist.org/). We calculated the proportion of threatened species by dividing the number of threatened species in each order by the total number of amphibian or bird species as appropriate. We did the same to find the proportion of amphibian and bird species in each order (i.e. dividing the number of species in each order by the total number of species).

## Results

There was substantial bias in the spatial distribution of evidence in conservation. Approximately 79% of amphibian studies and 76% of bird studies were conducted in either North America or Europe. Additionally, 56% of amphibian studies and 60% of bird studies were conducted in either the UK or USA. There were large spatial gaps in evidence in South America, Africa, Asia, and Russia for both amphibians and birds. There were few studies in the tropics or close to the poles (Figs.1 & 2).

**Figure 1.**
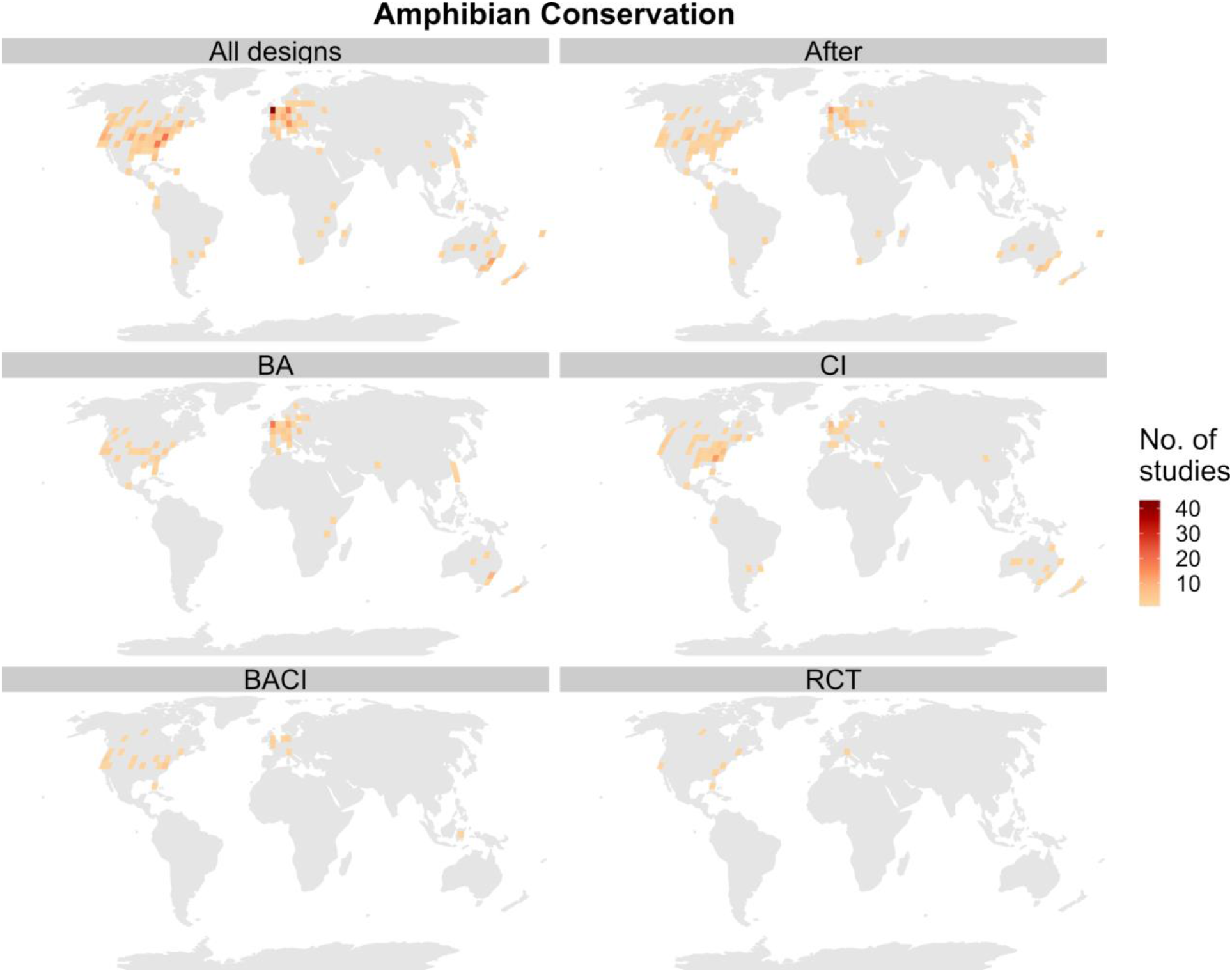
Spatial distribution of studies for amphibians using a Robinson projection and grid cells at a 4×4 degree resolution. Definitions of design acronyms are as follows: BA = Before-After; CI = Control-Impact; BACI = Before-After Control-Impact; RCT = Randomized Controlled Trial (see Table 1 for details of designs).

**Figure 2.**
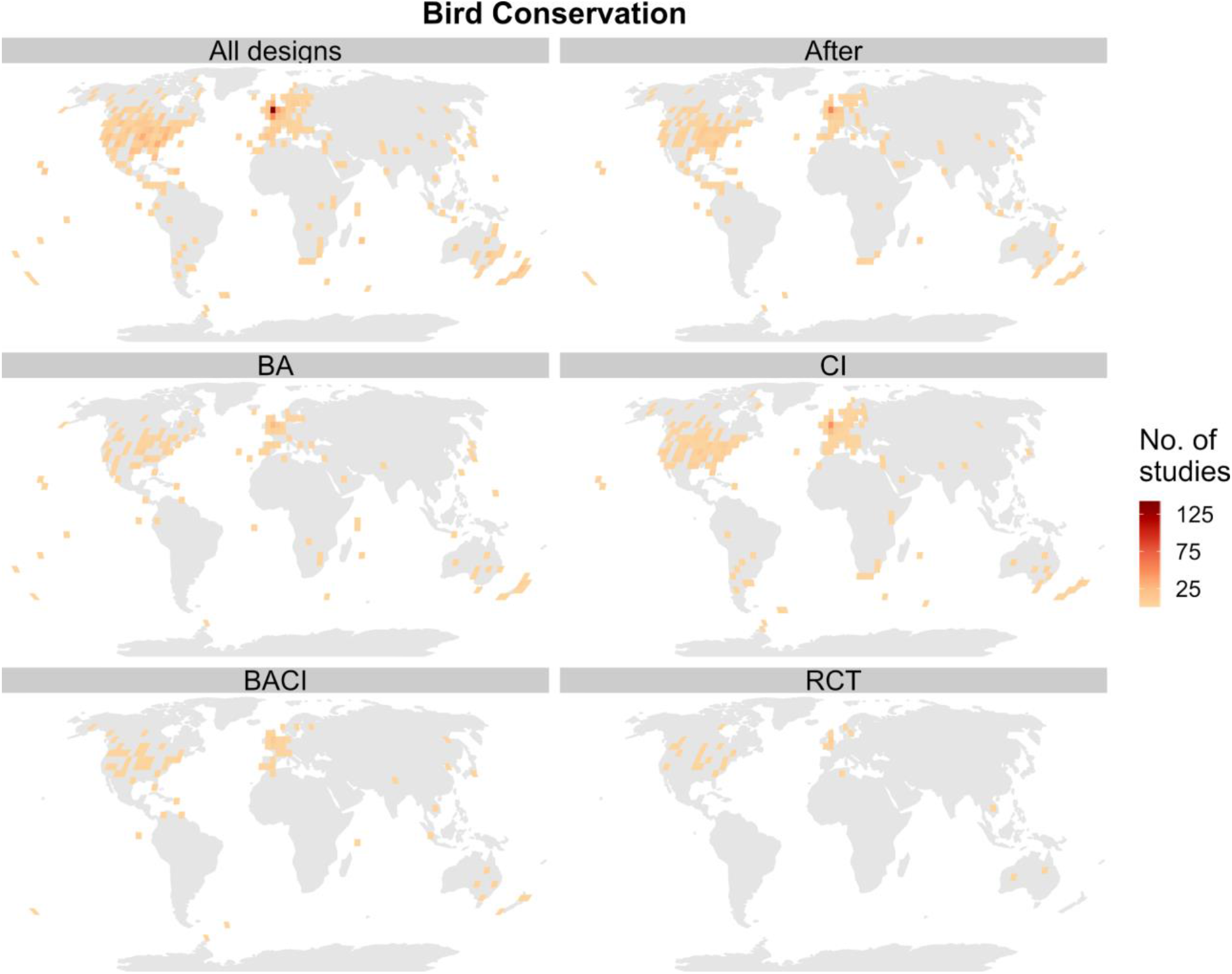
Spatial distribution of studies for birds using a Robinson projection and grid cells at a 4×4 degree resolution. Definitions of design acronyms are as follows: BA = Before-After; CI = Control-Impact; BACI = Before-After Control-Impact; RCT = Randomized Controlled Trial (see Table 1 for details of designs).

The geographical distribution of studies also depended on their design. Amphibian studies with the most robust study designs, BACI and RCT, were mainly concentrated in North America and Europe; these designs were almost absent from the tropics (Fig.1). BACI studies for amphibians were found on one more continent than RCTs (two versus three), but both only covered six countries (Table 2; Fig.1). BA studies for amphibians were found in 23 countries over five continents, whilst CI studies were found over six continents but only 18 countries. After studies for amphibians covered the greatest number of countries (31) across six continents (Table 2).

**Table 2.** Number of countries and continents where at least one study using a given study design was present (see Table 1 for details of designs).

|  | Amphibians |  | Birds |  |
| --- | --- | --- | --- | --- |
| Design | No. of countries | No. of continents | No. of countries | No. of continents |
| After | 31 | 6 | 46 | 7 |
| Control-Impact | 18 | 6 | 37 | 7 |
| Before-After | 23 | 5 | 37 | 7 |
| Before-After<br>Control-Impact | 6 | 3 | 20 | 7 |
| Randomized<br>Controlled Trial | 6 | 2 | 16 | 5 |

The evidence for birds had a greater geographical coverage than for amphibians, particularly in the tropics (Fig.2). RCT studies covered the fewest continents (5) compared with studies using other designs (7). RCT and BACI studies were also present in considerably fewer countries than After, CI and BA studies (Table 2).

The number of species, threatened species and data deficient species corresponded poorly with the number of studies in each 1×1 degree cell for both amphibians and birds - this was particularly pronounced for amphibians (Fig.3). Mutual information tests suggested there was almost no similarity between the geographical distributions of studies and either species, threatened species or data-deficient species (a value of zero suggests that the two distributions are independent of each other; mutual information values: all bird species = 0.009; threatened bird species = 0.004; data deficient bird species = 3×10^−5^; all amphibian species = 0.005; threatened amphibian species = 0.001; data deficient amphibian species = 4×10^−5^).

**Figure 3.**
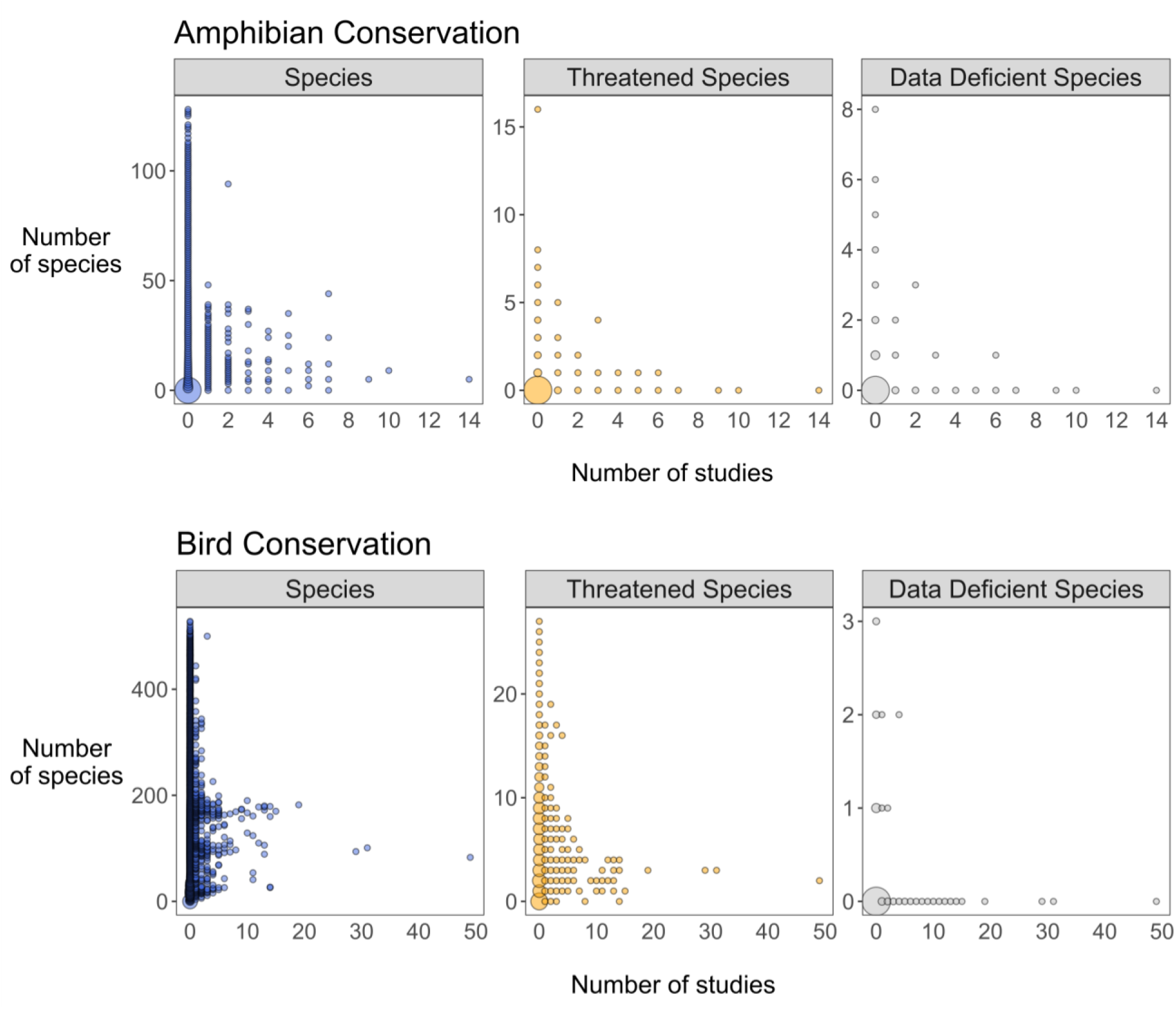
Comparison of the number of studies and the number of species (all species, threatened species and data deficient species) in 1×1 degree grid cells for amphibians and birds. The size of points is proportional to the number of points at those coordinates on the figure to aid visualization. Threatened species are those classified as vulnerable, endangered or critically endangered respectively, according to the IUCN Red List (IUCN 2019).

There was also substantial variation in the proportion of studies that tested interventions on different amphibian and bird orders relative to the proportion of species and threatened species each order contained. For birds, shorebirds (Charadriiformes), followed by waterfowl (Anseriformes) and falcons (Falconiformes) were better represented - i.e. high proportions of studies relative to proportions of threatened species (Fig.4). Songbirds (Passeriformes), parrots (Psittaciformes), pigeons (Columbiformes), and nightjars, hummingbirds and swifts (Caprimulgiformes), were the most underrepresented bird orders - i.e. low proportions of studies relative to threatened species. No studies were present for several bird orders, such as hornbills and hoopoes (Bucerotiformes; see red outlines to circles in Fig.4). For amphibians, frogs (Anura) were underrepresented, whilst salamanders (Caudata) were better represented relative to the proportion of threatened species. There was only a single test of the effectiveness of an intervention for the entire order of Caecilians (Gymnophiona; Fig.4). Patterns were different when considering the proportion of studies relative to the proportion of species in each bird order; most bird orders were relatively well represented apart from songbirds and orders for which we found no studies (Fig.S1). For amphibians, patterns in representation were similar for both the proportion of species and proportion of threatened species (Fig.S1).

**Figure 4.**
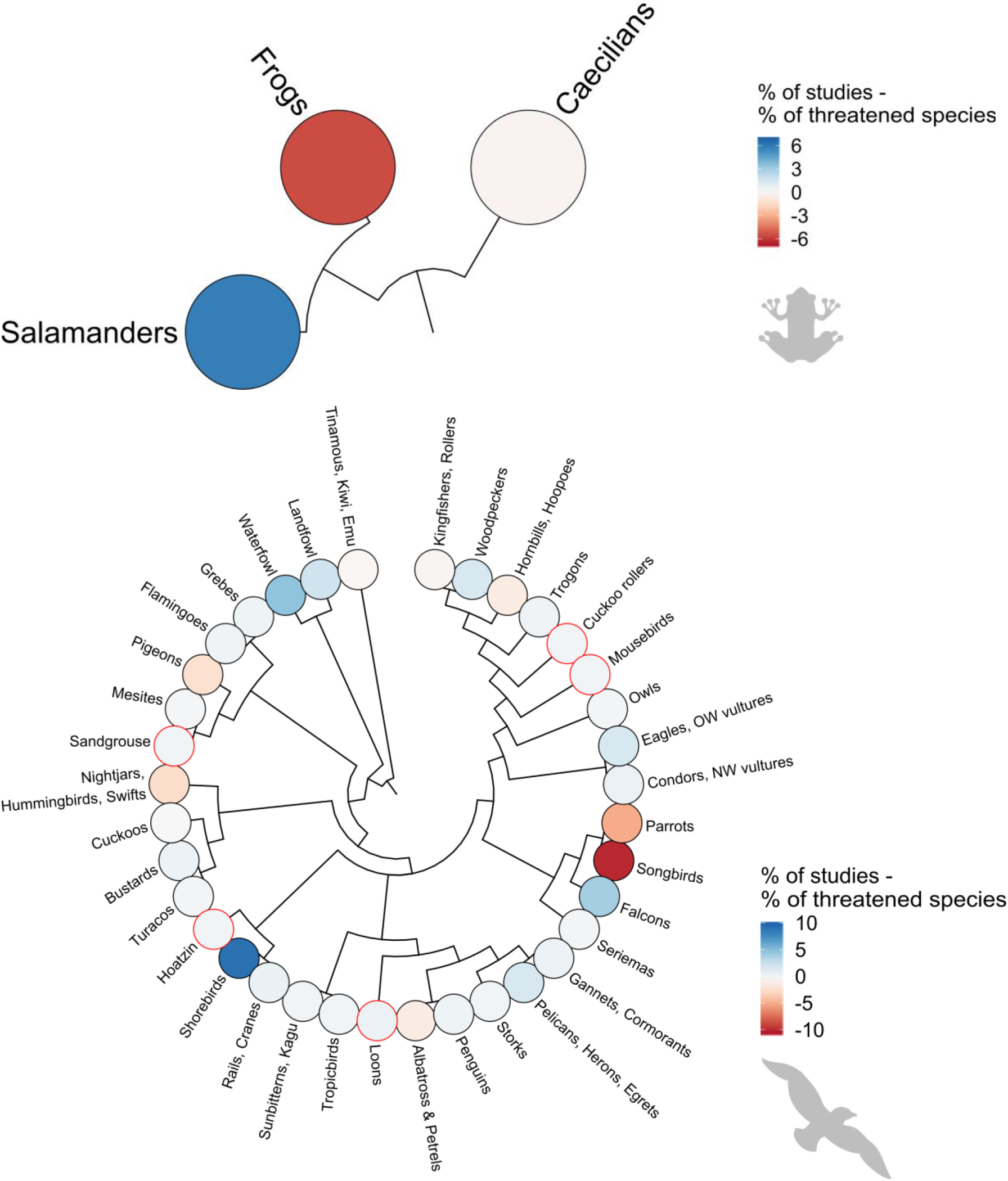
Percentage of studies minus percentage of threatened species in each order of amphibians and birds - percentages are relative to the total number of amphibian or bird studies and amphibian or bird species. Red outlines to circles indicate zero studies for that order. Darker blue colors indicate higher proportions of studies relative to the proportion of threatened species, whilst darker red colors indicate relatively lower proportions of studies.

We also found that the mean number of studies per intervention was low (less than three) for most biomes (Fig.5). The total number of interventions (containing at least one study) used to calculate the mean number of studies per intervention was 243 for birds and 74 for amphibians. For grassland and shrubland biomes there were fewer than one study per intervention for birds, but approximately 5.5 studies per intervention for amphibians. For forest biomes these patterns were reversed; there were approximately 5.7 studies per intervention for birds and approximately 2.8 studies per intervention for amphibians. The two biomes with most studies for both amphibians and birds were temperate broadleaf and mixed forests, and temperate grasslands, savannas and shrublands (Fig.5).

**Figure 5.**
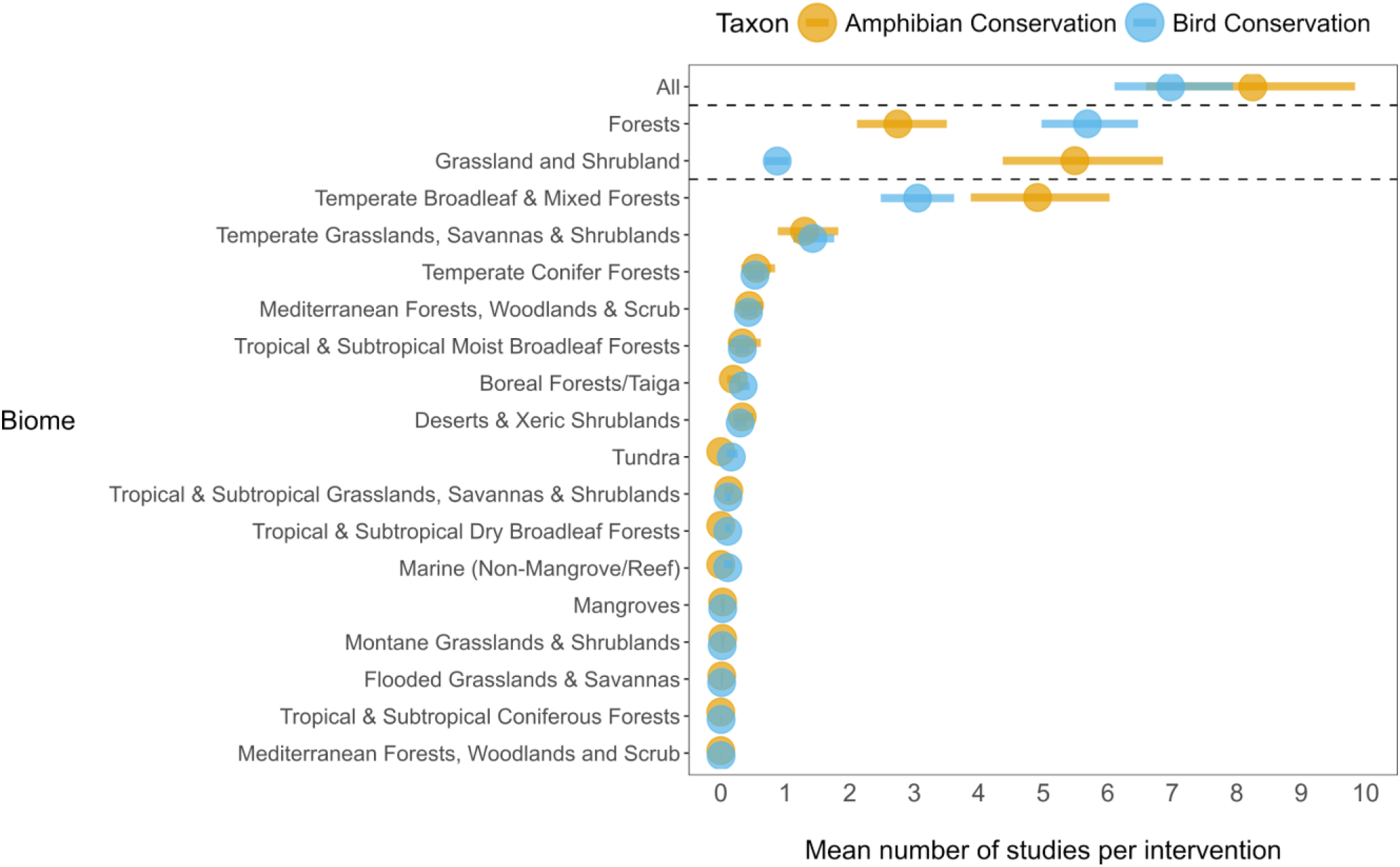
Mean number of studies per intervention for different biomes. Bars show bootstrapped 95% Confidence Intervals. The category ‘All’ refers to the mean number of studies per intervention in any biome, whilst the categories ‘Forests’ and ‘Grassland and Shrubland’ only include studies that were conducted in any forest or any grassland/shrubland biome, respectively.

The most commonly used metrics in amphibian conservation were mortality/survival (3.9 studies per intervention) and reproductive success (3.8 studies per intervention), whilst for birds, mortality/survival (3.9 studies per intervention) and abundance/density/cover (3.8 studies per intervention; Fig.6) were most common. On average, the effectiveness of each intervention was measured using 2.1 different metrics for amphibians and 3.3 metrics for birds.

**Figure 6.**
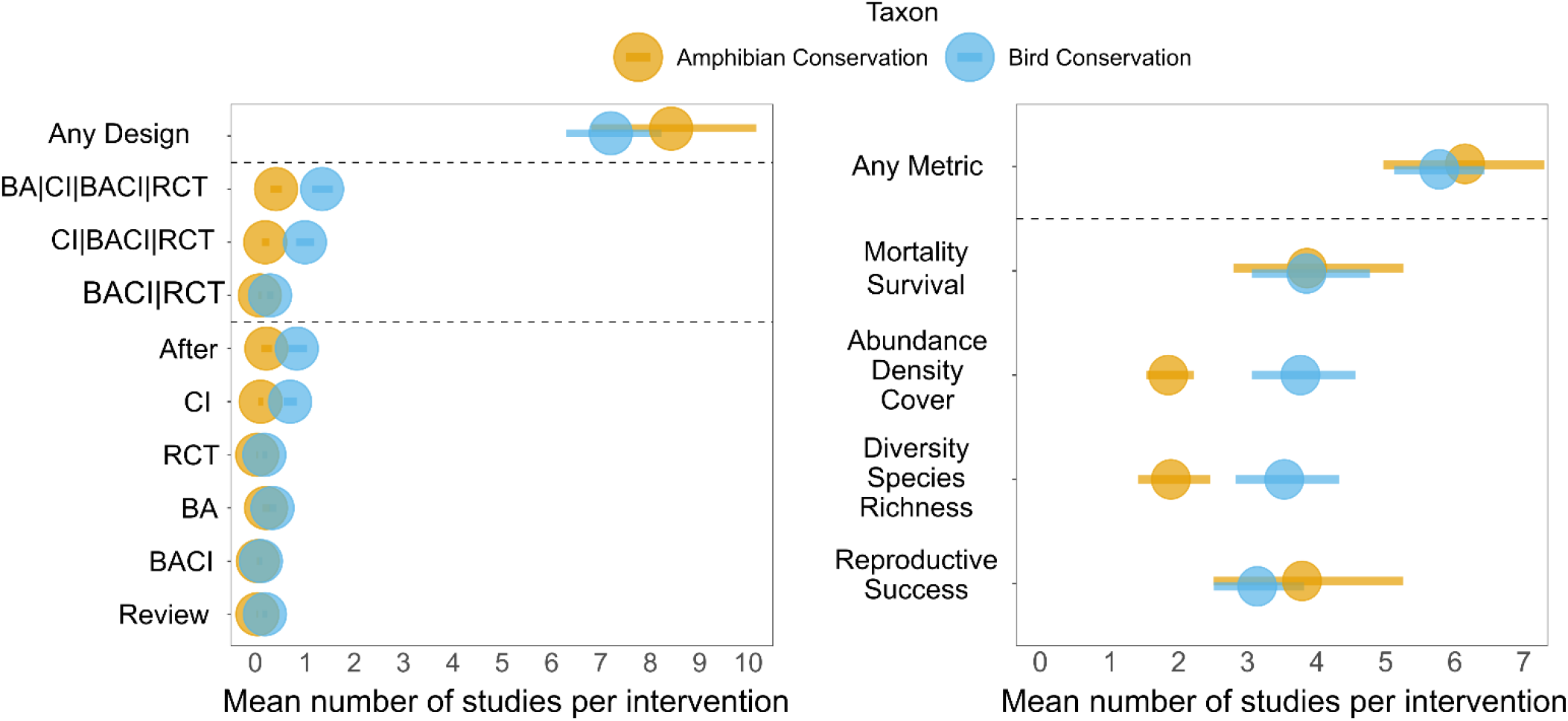
Mean number of studies per intervention with different designs and effectiveness metrics. ‘Any Metric’ refers to the mean number of studies using any of the four groups of metrics per intervention. ‘Any Design’ refers to the mean number of studies using any of the study designs per intervention. | symbolizes ‘or’ – i.e. BACI|RCT means studies with BACI or RCT designs. Bars show bootstrapped 95% Confidence Intervals.

Studies most commonly used the least robust After design, followed by CI and BA designs, for both amphibians and birds (Fig.6). There was a low number of studies per intervention when considering only studies with RCT, BACI, CI or BA designs (fewer than two for both amphibians and birds), and even fewer when we restricted the design type to RCT, BACI or CI and RCT or BACI designs (Fig.6).

## Discussion

Our work demonstrates that the evidence base for conservation suffers from severe geographical and taxonomic biases that may hamper our ability to make locally relevant evidence-based recommendations to decision-makers. Geographically, studies were concentrated in North America and Western Europe, whilst the number of studies in 1×1 degree cells showed no relationship with the number of species, and the number of threatened or data deficient species. Geographical bias was also indicated by the fact that the biomes with the highest mean number of studies per intervention were temperate broadleaf & mixed forests and temperate grasslands, savannas & shrublands. Taxonomically, certain orders were better studied relative to the number of threatened species they contained (e.g. salamanders for amphibians and shorebirds, falcons and waterfowl for birds), whilst some orders were not studied at all (e.g. sandgrouse and loons).

These results show similar latitudinal and geographic biases to other studies of the evidence base in conservation, such as Wilson et al. (2016), Di Marco et al. (2017), and Hickisch et al. (2019), but show different distributions of studies in the tropics to those from Burivalova et al. (2019). These differences probably result from our focus on all conservation interventions for amphibians and birds, whereas Burivalova et al. (2019) reviewed studies of social interventions to conserve entire habitats, such as payments for ecosystem services and community forest management. The paucity of evidence we found in the polar regions, Africa, Russia, the Middle East and South America broadly corresponds to patterns of publication density in Di Marco et al. (2017) and Hickisch et al. (2019). We also found that the United Kingdom rivalled the United States of America as a hotspot of evidence for these two taxonomic groups, which did not seem to be as apparent in Wilson et al. (2016) or Hickisch et al. (2019), but was in Di Marco et al. (2017). This hotspot contrasts, particularly for amphibians, with their low species richness in the UK (only seven native amphibian species). This is probably because we considered a different subset of studies, focusing only on studies that had quantitatively tested a conservation intervention, as opposed to any biodiversity or conservation-related publication. However, the Conservation Evidence database currently includes few studies from non-English language journals. Part of the geographic bias we found is likely attributable to the lack of studies from non-English language journals. As researchers conducting evidence synthesis, we must do more to seek out and collate evidence published in non-English languages - for example, Conservation Evidence has now searched over 150 non-English language journals, which will be included in future syntheses (Sutherland et al. 2019). This is particularly important given approximately 36% of the conservation literature is found in non-English language journals (Amano et al. 2016).

Some taxonomic groups were well represented in the literature relative to the total number of species and the number of threatened species in the group, while other taxonomic groups were very poorly represented (Fig.4, Figure S1) – as found in analyses of the wider conservation literature (Clark and May 2002; Fazey et al. 2005; Murray et al. 2015; Donaldson et al. 2016). An underrepresentation of threatened species is concerning because information on the effectiveness of interventions targeting threatened species is urgently required – particularly given substantial declines of bird fauna (Rosenberg et al. 2019) and severe threats to amphibians (Grant et al. 2019). While it can be challenging to design robust studies on rare species, where feasible, conservation scientists should prioritize testing the effectiveness of conservation interventions for threatened species. Equally, the absence of some orders from the literature testing conservation interventions is important because functional and ecological differences between taxonomic groups may make generalization of the effectiveness of interventions difficult. Investigating which interventions are likely to be effective in many local contexts is extremely important so we can prioritize the most important taxonomic gaps that need to be addressed in the evidence base for conservation.

Types of heterogeneity that may complicate the process of evidence synthesis were also present; for example, there was substantial variation in the number of studies with different study designs, and notably few studies used robust designs (e.g. RCT or BACI). This heterogeneity in study design was also linked to geographic biases, whereby studies with more robust designs (e.g. RCT or BACI) tended to be strongly concentrated in North America and Western Europe (particularly the United Kingdom) compared to less robust designs (e.g. BA, CI, After). This suggests that we do not only lack studies outside of North America and Western Europe, but also that the few studies that do exist outside these regions are likely to be of poor methodological quality and potentially biased (Christie et al. 2019). We should therefore prioritize future research effort on testing conservation interventions using robust study designs in these underrepresented regions and ensuring that they are published.

Amphibian and bird studies also used a variety of metrics to quantify the effectiveness of the same intervention - on average, approximately two metrics per intervention for amphibians and three for birds. Although using several metrics may help us better understand the overall effectiveness of an intervention, too many metrics can make evidence hard to synthesize, such as in systematic reviews and meta-analyses, and difficult to interpret for decision-makers. The diversity of metrics used could limit the number of studies that are directly comparable, hampering efforts to conduct meta-analyses and potentially making decision-making more difficult. This highlights the need for greater standardization of the metrics we use to assess conservation effectiveness (Mace & Baillie 2007; Capmourteres & Anand 2016; McQuatters-Gollop et al. 2019). Clearly, we are not advocating that we only use one standard type of metric in each study, but rather that we use the same set of metrics within each study to test a given conservation intervention - this will make studies that test the same intervention more directly comparable.

The gaps and biases we have highlighted in the literature on the effectiveness of conservation interventions represents a serious issue for the field of conservation. Although we were only able to analyze the literature up until 2012 for amphibians and 2011 for birds, we believe it is likely that these gaps and biases still persist to a greater or lesser extent in the evidence base for conservation. With limited resources we cannot afford to allocate research effort inefficiently. Therefore, the results of our work are extremely important for determining where future research effort on testing the effectiveness of conservation interventions should be invested. Future studies should target the poorly represented threatened taxa we identified, as well as poorly represented regions and biomes (e.g. the tropics). Although studies using more robust designs often require extra research effort, we also suggest some investment should be made to increase the use of robust designs outside of Western Europe and North America. Future studies should also use a consistent suite of metrics to ensure they present results that are relevant to a broader range of decision-makers and more directly comparable for evidence synthesis.

Future research is needed to identify specific priorities for taxonomic groups other than amphibians and birds, although the broad biases and gaps we identified in the conservation evidence literature are likely to apply to other taxonomic groups. We hope that by addressing geographic and taxonomic biases in the evidence base we can ensure more relevant evidence-based recommendations can be made to decision-makers. Similarly, addressing heterogeneity in the designs and metrics used by studies will hopefully allow evidence synthesis to be more efficient and effective. We believe that a more complete, robust, and standardized evidence base will enable conservation to become more evidence-based and, ultimately, more effective.

## Acknowledgements

We would like to thank Anne Mupepele for their useful comments on the manuscript and all past and present members of the Conservation Evidence project.

## Author funding sources

TA was supported by the Grantham Foundation for the Protection of the Environment, the Kenneth Miller Trust and the Australian Research Council Future Fellowship (FT180100354); WJS, PAM, CFRW, SOP and GES are supported by Arcadia and The David and Claudia Harding Foundation; BIS and APC were supported by the Natural Environment Research Council as part of the Cambridge Earth System Science NERC DTP [NE/L002507/1]. BIS is also supported by the Natural Environment Research Council [NE/S001395/1].

