## Appendices S1-S4 for "The challenge of heterogeneous evidence in conservation": Appendix_S3_anon.R

```
debug(utils:::unpackPkgZip)
require("RCurl")
require('tm')
library(doParallel)
require(XML)

master = matrix(nrow=10000,ncol=5)
nstud = 9000


masterloop <- function(zeta){
  z=zeta
  st2835 = getURL(paste("https://www.conservationevidence.com/individual-study/", z,sep=""),ssl.verifypeer=FALSE) ### access conservation evidence website and summary of study with ID zeta (e.g. 1, 2, 3 ...)

  st2835tree = htmlTreeParse(st2835,useInternal = TRUE)
  
  st2835tree.html=do.call(paste, as.list(capture.output(st2835tree)))

  masterlog = NULL
  if(length(grep("<h2>Summary</h2>",st2835tree.html))==0){ ### Only look at website pages that do not have an old-style heading, these are archived 
                                                            ### and not used anymore and are not assgined to an intervention or synopsis
    
    
    st2835tree.parse = unlist(xpathApply(st2835tree,path="//p",fun=xmlValue))

    
    if (length(grep("This option allows you to download this individual study.",st2835tree.parse))==0){
      st2835.txt = list()
      j=1
      for (i in 4:(length(st2835tree.parse)-3)) {
        st2835.txt[[j]] = tolower(as.character(st2835tree.parse[i]))
        j=j+1
      }
    }
    
    
    if (length(grep("This option allows you to download this individual study.",st2835tree.parse))>0){
      st2835.txt = list()
      j=1
      for (i in 4:(max(grep("This option allows you to download this individual study.",st2835tree.parse))-1)) {
        st2835.txt[[j]] =     tolower(as.character(st2835tree.parse[i]))
        j=j+1
      }
    }
    
    
    
    masterlog = matrix(ncol=10,nrow=length(st2835.txt))
    
    for(i in 1:length(st2835.txt)){
      
      ###density
      if(length(grep("/plot",st2835.txt[[i]])) > 0 |length(grep("volume of arthropods",st2835.txt[[i]])) > 0 |length(grep("earthworms/m2",st2835.txt[[i]])) > 0 |length(grep(" per fish",st2835.txt[[i]])) > 0 |length(grep("density",st2835.txt[[i]])) > 0 |length(grep("densities",st2835.txt[[i]])) > 0 |length(grep("captures increased",st2835.txt[[i]])) > 0|length(grep("captures decreased",st2835.txt[[i]])) > 0){masterlog[i,1]=1}
      if(length(grep("/plot",st2835.txt[[i]])) == 0 &length(grep("volume of arthropods",st2835.txt[[i]])) == 0 &length(grep("earthworms/m2",st2835.txt[[i]])) == 0 &length(grep(" per fish",st2835.txt[[i]])) == 0 &length(grep("density",st2835.txt[[i]])) == 0 &length(grep("densities",st2835.txt[[i]])) == 0 &length(grep("captures increased",st2835.txt[[i]])) == 0&length(grep("captures decreased",st2835.txt[[i]])) == 0){masterlog[i,1]=0}
      if(length(grep("effect of.{,20}density",st2835.txt[[i]])) > 0 | length(grep("effect of.{,20}density",st2835.txt[[i]])) > 0){masterlog[i,1]=0}
      
      ##abundance
      if(length(grep("were counted",st2835.txt[[i]])) > 0 |length(grep("more nest-searching",st2835.txt[[i]])) > 0 |length(grep("more foraging",st2835.txt[[i]])) > 0 |length(grep("number of breeding",st2835.txt[[i]])) > 0 |length(grep("more microorganisms",st2835.txt[[i]])) > 0 |length(grep("similar numbers",st2835.txt[[i]])) > 0 |length(grep("more numerous",st2835.txt[[i]])) > 0 |length(grep("numbers of nest-searching",st2835.txt[[i]])) > 0 |length(grep("numbers of foraging",st2835.txt[[i]])) > 0 |length(grep("pest numbers",st2835.txt[[i]])) > 0 |length(grep("doubled a population",st2835.txt[[i]])) > 0 |length(grep("population.{,30}doubl",st2835.txt[[i]])) > 0 |length(grep(" more pollinators ",st2835.txt[[i]])) > 0 |length(grep(" more.{,25}were seen ",st2835.txt[[i]])) > 0 |length(grep("counts of",st2835.txt[[i]])) > 0 |length(grep("total number",st2835.txt[[i]])) > 0 |length(grep("abundance",st2835.txt[[i]])) > 0 |length(grep("more abundant",st2835.txt[[i]])) > 0 |length(grep("less abundant",st2835.txt[[i]])) > 0 |length(grep("population.{,50}increase",st2835.txt[[i]])) > 0 |length(grep("population.{,50}decrease",st2835.txt[[i]])) > 0 |length(grep("in.{,25}number",st2835.txt[[i]])) > 0|length(grep("crease in.{,35}population",st2835.txt[[i]])) > 0|length(grep("numbers.{,20}..creased",st2835.txt[[i]])) > 0|length(grep("higher.{,20}number",st2835.txt[[i]])) > 0|length(grep("lower.{,20}number",st2835.txt[[i]])) > 0){masterlog[i,2]=1}
      if(length(grep("were counted",st2835.txt[[i]])) == 0 &length(grep("more nest-searching",st2835.txt[[i]])) == 0 &length(grep("more foraging",st2835.txt[[i]])) == 0 &length(grep("number of breeding",st2835.txt[[i]])) == 0 &length(grep("more microorganisms",st2835.txt[[i]])) == 0 &length(grep("similar numbers",st2835.txt[[i]])) == 0 &length(grep("more numerous",st2835.txt[[i]])) == 0 &length(grep("numbers of nest-searching",st2835.txt[[i]])) == 0 &length(grep("numbers of foraging",st2835.txt[[i]])) == 0 &length(grep("pest numbers",st2835.txt[[i]])) == 0 &length(grep("doubled a population",st2835.txt[[i]])) == 0 &length(grep("population.{,30}doubl",st2835.txt[[i]])) == 0 &length(grep(" more pollinators ",st2835.txt[[i]])) == 0 &length(grep(" more.{,25}were seen ",st2835.txt[[i]])) == 0 &length(grep("counts of",st2835.txt[[i]])) == 0 &length(grep("total number",st2835.txt[[i]])) == 0&length(grep("abundance",st2835.txt[[i]])) == 0 & length(grep("more abundant",st2835.txt[[i]])) == 0 &length(grep("less abundant",st2835.txt[[i]])) == 0 &length(grep("population.{,50}increase",st2835.txt[[i]])) == 0 &length(grep("population.{,50}decrease",st2835.txt[[i]])) == 0&length(grep("crease in.{,35}population",st2835.txt[[i]])) == 0&length(grep("numbers.{,20}..creased",st2835.txt[[i]])) == 0&length(grep("higher.{,20}number",st2835.txt[[i]])) == 0&length(grep("lower.{,20}number",st2835.txt[[i]])) == 0){masterlog[i,2]=0}
      
      ##diversity
      if(length(grep(" more.{,25}species ",st2835.txt[[i]])) > 0 |length(grep("richness",st2835.txt[[i]])) > 0 |length(grep(" diversity ",st2835.txt[[i]])) > 0|length(grep(" numbers of species ",st2835.txt[[i]])) > 0 |length(grep(" number of species ",st2835.txt[[i]])) > 0|length(grep(" numbers of.{,25}species ",st2835.txt[[i]])) > 0 |length(grep(" number of.{,25}species ",st2835.txt[[i]])) > 0 |length(grep("community composition",st2835.txt[[i]])) > 0|length(grep("species composition",st2835.txt[[i]])) > 0){masterlog[i,3]=1}
      if(length(grep(" more.{,25}species ",st2835.txt[[i]])) == 0 &length(grep("richness",st2835.txt[[i]])) == 0 &length(grep(" diversity ",st2835.txt[[i]])) == 0&length(grep(" numbers of species ",st2835.txt[[i]])) == 0&length(grep(" number of species ",st2835.txt[[i]])) == 0&length(grep(" numbers of.{,25}species ",st2835.txt[[i]])) == 0&length(grep(" number of.{,25}species ",st2835.txt[[i]])) == 0&length(grep("community composition",st2835.txt[[i]])) == 0&length(grep("species composition",st2835.txt[[i]])) == 0){masterlog[i,3]=0}
      if(length(grep(" small numbers of these species ",st2835.txt[[i]])) > 0){masterlog[i,3]=0}
      
      ##cover
      if(length(grep("effect on.{,15}cover",st2835.txt[[i]])) > 0 |length(grep("herbaceous cover",st2835.txt[[i]])) > 0 |length(grep("percentage.{,15}cover",st2835.txt[[i]])) > 0 |length(grep("% cover",st2835.txt[[i]])) > 0 |length(grep("average.{,15}cover",st2835.txt[[i]])) > 0|length(grep("species cover",st2835.txt[[i]])) > 0|length(grep("greater.{,15}cover",st2835.txt[[i]])) > 0|length(grep("lower.{,15}cover",st2835.txt[[i]])) > 0|length(grep("higher.{,15}cover ",st2835.txt[[i]])) > 0|length(grep("less.{,15}cover",st2835.txt[[i]])) > 0|length(grep("more.{,15}cover",st2835.txt[[i]])) > 0|length(grep(" cover of",st2835.txt[[i]]))>0|length(grep(" cover decreased",st2835.txt[[i]])) > 0|length(grep(" cover increased",st2835.txt[[i]])) > 0|length(grep("decreased.{,15}cover",st2835.txt[[i]])) > 0|length(grep("..creased.{,15}cover",st2835.txt[[i]])) > 0|length(grep("less.{,25}cover",st2835.txt[[i]])) > 0|length(grep("more.{,25}cover",st2835.txt[[i]])) > 0 |length(grep("total.{,25}cover",st2835.txt[[i]])) > 0 | length(grep("proportion of.{,15}cover",st2835.txt[[i]])) > 0 | length(grep("cover was higher",st2835.txt[[i]])) > 0 | length(grep("cover was lower",st2835.txt[[i]])) > 0 | length(grep("% of the surface was cover",st2835.txt[[i]])) > 0 ){masterlog[i,4]=1}
      if(length(grep("effect on.{,15}cover",st2835.txt[[i]])) == 0 &length(grep("herbaceous cover",st2835.txt[[i]])) == 0 &length(grep("percentage.{,15}cover",st2835.txt[[i]])) == 0 &length(grep("% cover",st2835.txt[[i]])) == 0 &length(grep("average.{,15}cover",st2835.txt[[i]])) == 0&length(grep("species cover",st2835.txt[[i]])) == 0&length(grep("greater.{,15}cover",st2835.txt[[i]])) == 0&length(grep("lower.{,15}cover",st2835.txt[[i]])) == 0&length(grep("higher.{,15}cover ",st2835.txt[[i]])) == 0&length(grep("less.{,15}cover",st2835.txt[[i]])) == 0&length(grep("more.{,15}cover",st2835.txt[[i]])) == 0&length(grep(" cover of",st2835.txt[[i]])) == 0&length(grep(" cover decreased",st2835.txt[[i]])) == 0&length(grep(" cover increased",st2835.txt[[i]])) == 0&length(grep("decreased.{,15}cover",st2835.txt[[i]])) == 0&length(grep("..creased.{,15}cover",st2835.txt[[i]])) == 0&length(grep("less.{,25}cover",st2835.txt[[i]])) == 0&length(grep("more.{,25}cover",st2835.txt[[i]])) == 0 &length(grep("total.{,25}cover",st2835.txt[[i]])) == 0 &length(grep("proportion of.{,15}cover",st2835.txt[[i]])) == 0 &length(grep("cover was higher",st2835.txt[[i]])) == 0&length(grep("cover was lower",st2835.txt[[i]])) == 0&length(grep("% of the surface was cover",st2835.txt[[i]])) == 0){masterlog[i,4]=0}
      if(length(grep(" cover plot",st2835.txt[[i]])) > 0){masterlog[i,4]=0}

      #####survival
      if(length(grep("surviv..",st2835.txt[[i]])) > 0 ){masterlog[i,5]=1}
      if(length(grep("surviv..",st2835.txt[[i]])) == 0 ){masterlog[i,5]=0}
      
      #####reproductive success
      if(length(grep("breeding pairs ..creased",st2835.txt[[i]])) > 0 |length(grep("fledg",st2835.txt[[i]])) > 0 |length(grep("brood",st2835.txt[[i]])) > 0 |length(grep("% of egg",st2835.txt[[i]])) > 0 |length(grep("nestlings produced",st2835.txt[[i]])) > 0 |length(grep("proportion of females with chicks",st2835.txt[[i]])) > 0 |length(grep(" success ",st2835.txt[[i]])) > 0 | length(grep("clutch",st2835.txt[[i]])) > 0 | length(grep("significantly fewer (.*?) nests",st2835.txt[[i]])) > 0| length(grep("significantly less (.*?) nests",st2835.txt[[i]])) > 0| length(grep("significantly more (.*?) nests",st2835.txt[[i]])) > 0| length(grep("more nests",st2835.txt[[i]])) > 0 | length(grep("less nests",st2835.txt[[i]])) > 0 | length(grep("fewer nests",st2835.txt[[i]])) > 0){masterlog[i,6]=1}
      if(length(grep("breeding pairs ..creased",st2835.txt[[i]])) == 0 &length(grep("fledg",st2835.txt[[i]])) == 0 &length(grep("brood",st2835.txt[[i]])) == 0 &length(grep("% of egg",st2835.txt[[i]])) == 0 &length(grep("nestlings produced",st2835.txt[[i]])) == 0 &length(grep("proportion of females with chicks",st2835.txt[[i]])) == 0 &length(grep(" success ",st2835.txt[[i]])) == 0 & length(grep("clutch",st2835.txt[[i]])) == 0& length(grep("significantly fewer (.*?) nests",st2835.txt[[i]])) == 0& length(grep("significantly less (.*?) nests",st2835.txt[[i]])) == 0& length(grep("significantly more (.*?) nests",st2835.txt[[i]])) == 0& length(grep("more nests",st2835.txt[[i]])) == 0 & length(grep("less nests",st2835.txt[[i]])) == 0 & length(grep("fewer nests",st2835.txt[[i]])) == 0){masterlog[i,6]=0}
      
      #####mortality
      if(length(grep("% predation",st2835.txt[[i]])) > 0 |length(grep("mortality",st2835.txt[[i]])) > 0 |length(grep("predation rate",st2835.txt[[i]])) > 0 ){masterlog[i,7]=1}
      if(length(grep("% predation",st2835.txt[[i]])) == 0 &length(grep("mortality",st2835.txt[[i]])) == 0 &length(grep("predation rate",st2835.txt[[i]])) == 0 ){masterlog[i,7]=0}

     
      ###Error
      if(length(grep("url was not found",st2835.txt[[i]])) > 0 ){masterlog[i,]=rep(NA,NCOL(masterlog))}
      if(length(grep("server error",st2835.txt[[i]])) > 0 ){masterlog[i,]=rep(NA,NCOL(masterlog))}
      if(length(grep("summarised by",st2835.txt[[i]])) > 0 ){masterlog[i,]=rep(NA,NCOL(masterlog))}
      if(length(grep("no summary",st2835.txt[[i]])) > 0 ){masterlog[i,]=rep(NA,NCOL(masterlog))}
      
      
      ###blank text
      if(length(grep("t",st2835.txt[[i]])) == 0 ){masterlog[i,]=rep(NA,NCOL(masterlog))}
      if(length(grep("laboratory",st2835.txt[[i]])) == 0 &length(grep("comparison",st2835.txt[[i]])) == 0 &length(grep("study",st2835.txt[[i]])) == 0 & length(grep("experiment",st2835.txt[[i]])) == 0 & length(grep("trial",st2835.txt[[i]])) == 0 ){masterlog[i,]=rep(NA,NCOL(masterlog))}
      
      masterlog[,8] = z

    }
    
    masterlog = masterlog[which(is.na(masterlog[,1])==FALSE),]
    synops = regmatches(st2835tree.html, gregexpr('(?<=alt=\").*?(?=\">)', st2835tree.html, perl=T))
    if(length(synops[[1]]) == 0) {synops2 = NA}
    if(length(synops[[1]]) == 1) {synops2 = synops[[1]]}
    if(length(synops[[1]]) > 1 ) {synops2 = synops[[1]][1:(length(synops[[1]])-1)]}
    if(is.null(nrow(masterlog))==TRUE){
      masterlog[9] = synops2[1]
    }
    
    
    interv = unlist(c(regmatches(st2835tree.html, gregexpr('(?<=actions/).*?(?=</a>)', st2835tree.html, perl=T))))
    interv = gsub('\">','',interv)
    interv = gsub('\\d','',interv)
    
    if(length(interv) == 0) {interv2 = NA}
    if(length(interv) == 1) {interv2 = interv}
    if(length(interv) > 1 ) {interv2 = interv[1:(length(interv))]}
    if(is.null(nrow(masterlog))==TRUE){
      masterlog[10] = interv2[1]
    }
    
    if(is.null(nrow(masterlog))==FALSE){
      masterlog[,9] = synops2[1:nrow(masterlog)]
      masterlog[,10] = interv2[1:nrow(masterlog)]
      if(length(which(duplicated(masterlog)==TRUE))>0){
        masterlog = masterlog[-which(duplicated(masterlog)==TRUE),]
      }
      
    }
    
    if(NROW(masterlog)>1 & NCOL(masterlog)>1){
      for(o in 1:NROW(masterlog)){
        if(is.na(masterlog[o,1])==TRUE){masterlog[o,]=rep(NA,NCOL(masterlog))}
      }
    }
    
    if(NROW(masterlog)<2 | NCOL(masterlog)<2){
      if(is.na(masterlog[1])==TRUE){masterlog=rep(NA,length(masterlog))}
    }
  }
  if(length(masterlog)==0){ masterlog = c(rep(NA,7),z,rep(NA,2))}
  
  return(masterlog)
}


#####Check function in serial
samp=sample(1:1698,1)
masterloop(samp)

#### Run in parallel

library(doParallel)
y = c(1:1697,1699:nstud) #webpage 1698 causes an error and contains and old study not assigned to a synopsis

c1 <- makeCluster(8)

###tell cluster what packages it needs too
clusterEvalQ(c1,c(library(RCurl),library(tm),library(XML)))

master <-  parSapplyLB(c1,y,masterloop)

stopCluster(c1)

master1 = do.call(rbind,master)

str(master)
str(master1)

colnames(master1)=c("density","abundance","diversity/richness","cover","survival","reprosuccess","mortality","pageIDt","synopsis","intervention")
head(master1)
```
