## Supplementary material for "The challenge of heterogeneous evidence in conservation": Figure S1

**Supporting Information**

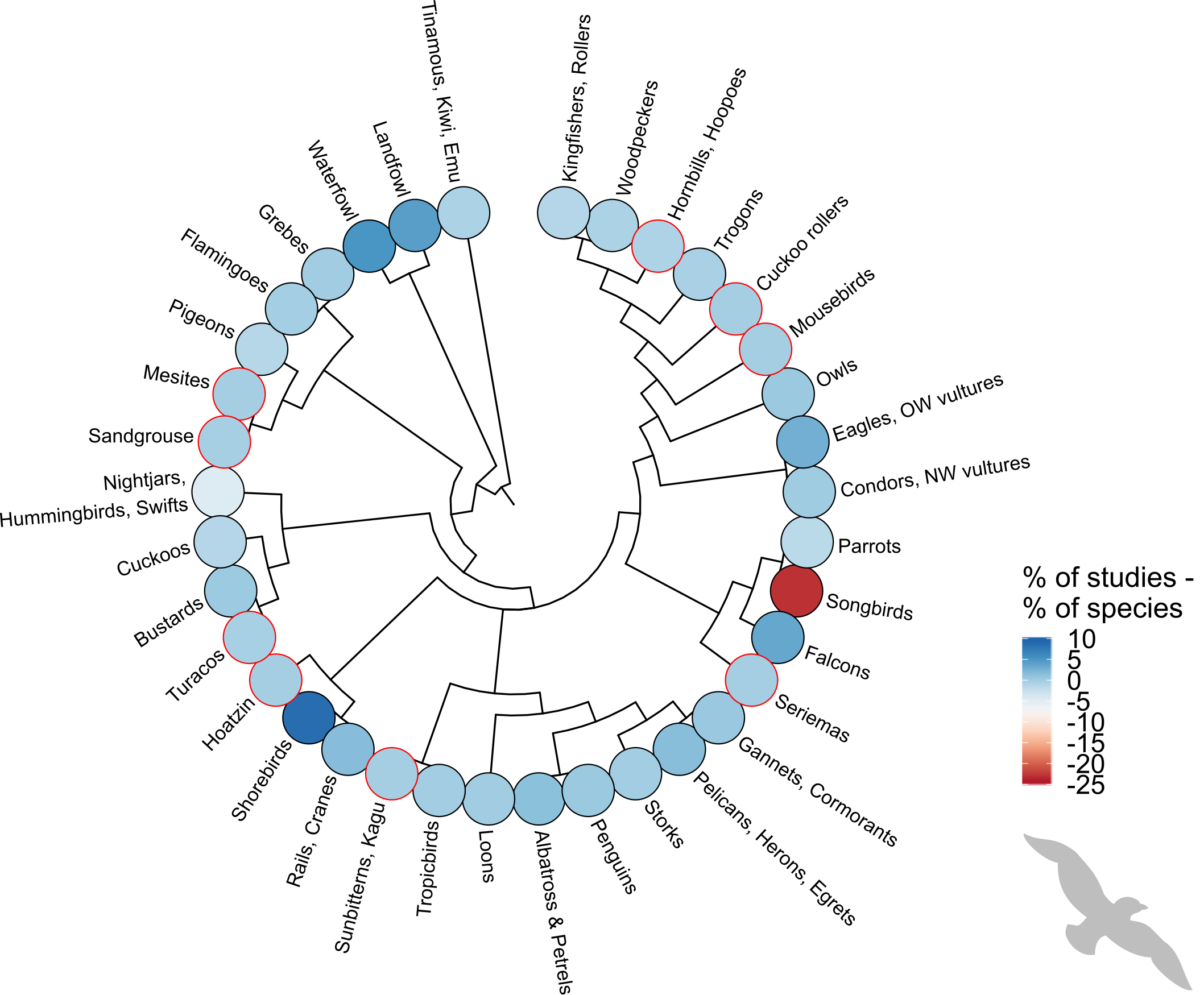

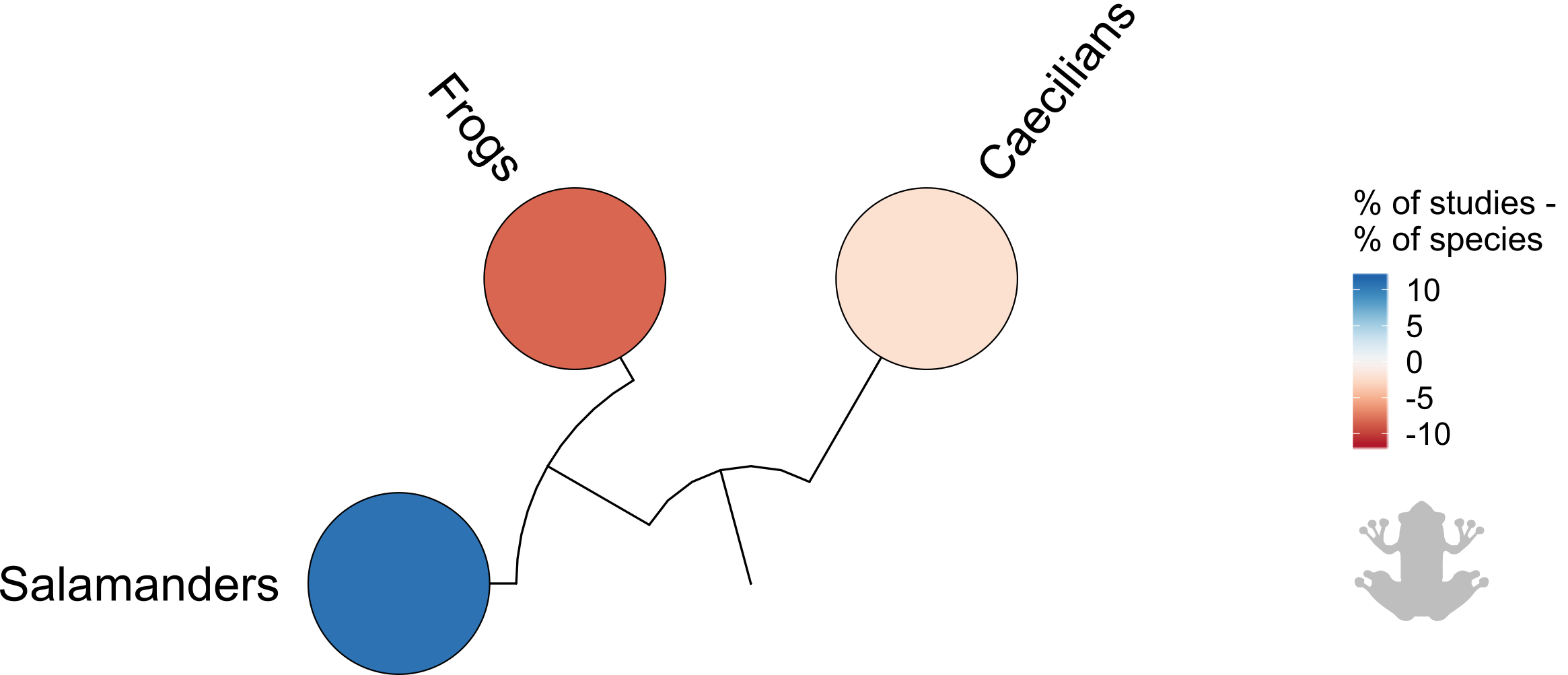

Figure S1 - Percentage of studies minus percentage of species in each order of amphibians and birds - percentages are relative to the total number of amphibian or bird studies and amphibian or bird species. Red outlines to circles indicate zero studies for that order. Darker blue colors indicate higher proportions of studies relative to the proportion of species, whilst darker red colors indicate relatively lower proportions of studies.
